# Prokaryotic community shifts during soil formation on sands in the tundra zone

**DOI:** 10.1101/449926

**Authors:** Alena Zhelezova, Timofey Chernov, Azida Tkhakakhova, Natalya Xenofontova, Mikhail Semenov, Olga Kutovaya

## Abstract

A chronosequence approach, *i.e.*, a comparison of spatially distinct plots with different stages of succession, is commonly used for studying microbial community dynamics during paedogenesis. The successional traits of prokaryotic communities following sand fixation processes have previously been characterized for arid and semi-arid regions, but they have not been considered for the tundra zone, where the environmental conditions are unfavourable for the establishment of complicated biocoenoses. In this research, we characterized the prokaryotic diversity and abundance of microbial genes found in a typical tundra and wooded tundra along a gradient of increasing vegetation – unfixed aeolian sand, semi-fixed surfaces with mosses and lichens, and mature soil under fully developed plant cover. Microbial communities from typical tundra and wooded tundra plots at three stages of sand fixation were compared using quantitative polymerase chain reaction (qPCR) and high-throughput sequencing of 16S rRNA gene libraries. The abundances of ribosomal genes increased gradually in both chronosequences, and a similar trend was observed for the functional genes related to the nitrogen cycle (*nifH*, bacterial *amoA, nirK* and *nirS*). The relative abundance of *Planctomycetes* increased, while those of *Thaumarchaeota, Cyanobacteria* and *Chloroflexi* decreased from unfixed sands to mature soils. According to β-diversity analysis, prokaryotic communities of unfixed sands were more heterogeneous compared to those of mature soils. Despite the differences in the plant cover of the two mature soils, the structural compositions of the prokaryotic communities were shaped in the same way.

## Introduction

For the investigation of microbial succession during soil-forming processes, a chronosequence approach, i.e., a comparison of spatially distinct plots of different ages, is commonly used. Currently, chronosequences of soil formation can be observed in areas with variable climatic conditions and on omnigenous parent material, such as glacial retreats [1–4], sand dunes [5–7], volcanic rocks [8], or anthropogenic landscapes [9]. Changes in microbial community structure during the process of sand dune fixation have mostly been studied for arid and semi-arid regions [6,10,11] and for coastal environments [5]. Currently, successional traits during sand fixation in the cold climate of the tundra have received increasing attention. The environmental conditions for soil formation in the tundra zone are specific, when the sandy substrates are depleted of nutrients, and the average temperature is unfavourable for complicated biogeocoenoses.

While the succession of plant communities is relatively well studied, information on the prokaryotic community assemblage during soil formation is still lacking. It is known that the first organisms to colonize parent rock are phototrophs, diazotrophs, chemolithotrophs and heterotrophs, whose taxonomic composition depends on the substrate properties [4,8]. Several bacterial phyla have been suggested to be associated with the initial stages of soil formation, mainly *Bacteroidetes* [3] and *Cyanobacteria* [12,13]. The prokaryotic community acts as the primary producer of organic matter and modifies the parent material for further colonization by plants. Available nitrogen is a limiting factor of plant growth, especially on lean substrates in cold environments [14–16]. Prokaryotes are able to perform nitrogen fixation, which leads to the accumulation of available nitrogen during inhabitation of barren substrates, such as rocks and sands [4]. Both archaeal and bacterial ammonia oxidizers produce nitrate (NO_3_^−^), which appears to be a crucial form of nitrogen for plants in the tundra zone [17]. Denitrification is a multi-step process of full or partial NO_3_^−^ reduction, which may lead to nitrogen losses through N_2_ and N_2_O emission [18,19]. Additionally, the presence of vegetation shapes prokaryotic community structure during soil formation [20]. In comparison to bacteria, fungi are less adapted to life on barren substrates and depend strongly on plants during the early stages of colonization [21].

The diversity of soil microorganisms changes during the process of soil formation; however, there is no distinct and universal pattern of prokaryotic diversity shifts with the successional stage of paedogenesis [2,3,20,22]. Previous studies have shown that at the earliest stages of soil formation after the retreat of glaciers (0 - 100 years), the bacterial diversity was relatively low, whereas it increased with the age of soil [2] or was the highest in middle-aged soils [3]. In contrast, in a longer timescale of ecosystem development (60 - 120 000 years), the diversity of the soil prokaryotic community decreased with the site age [22]. The patterns of prokaryotic diversity change among chronosequences of soil formation have been mostly studied for glacier retreats but not for soils formed on aeolian sand dunes.

The aim of this research was to reveal the traits of microbial community succession during sand fixation in the tundra zone. Two chronosequences of soil formation on aeolian sands with similar initial stages and different mature vegetation (typical tundra and wooded tundra) were compared. Taxonomic composition and diversity of the prokaryotic community, the abundances of bacterial, archaeal, and fungal ribosomal genes and functional genes related to the N cycle were estimated for three stages of sand fixation (unfixed sand - semi-fixed surface - mature soil). We hypothesized that 1) the abundance of prokaryotic communities increases among the chronosequences, 2) α-diversity of prokaryotic communities increases gradually with soil formation and plant colonization on sands, and 3) unfixed sands harbour similar prokaryotic community structures, while the communities in mature soils under the two vegetation types vary from each other.

## Materials and methods

### Sampling site description

Sand fixation chronosequences at two sites on the shores of the Pechora River (Northwestern Russia, Nenetsia region) were studied. This region is located in the southern tundra zone with a humid subarctic climate and an average annual temperature of −3.6 °C. The mean annual precipitation is 445 mm. For both sites, sampling was performed in August 2015 on three types of surfaces: 1 – unfixed aeolian sand, 2 – semi-fixed surface with mosses and lichens, and 3 – mature soil under developed plant cover (Fig 1). The two sites differed in the plant cover that developed on mature soil - typical tundra vegetation with subshrubs (Site I) and wooded tundra with rare trees and subshrubs (Site II).

**Fig. 1.**
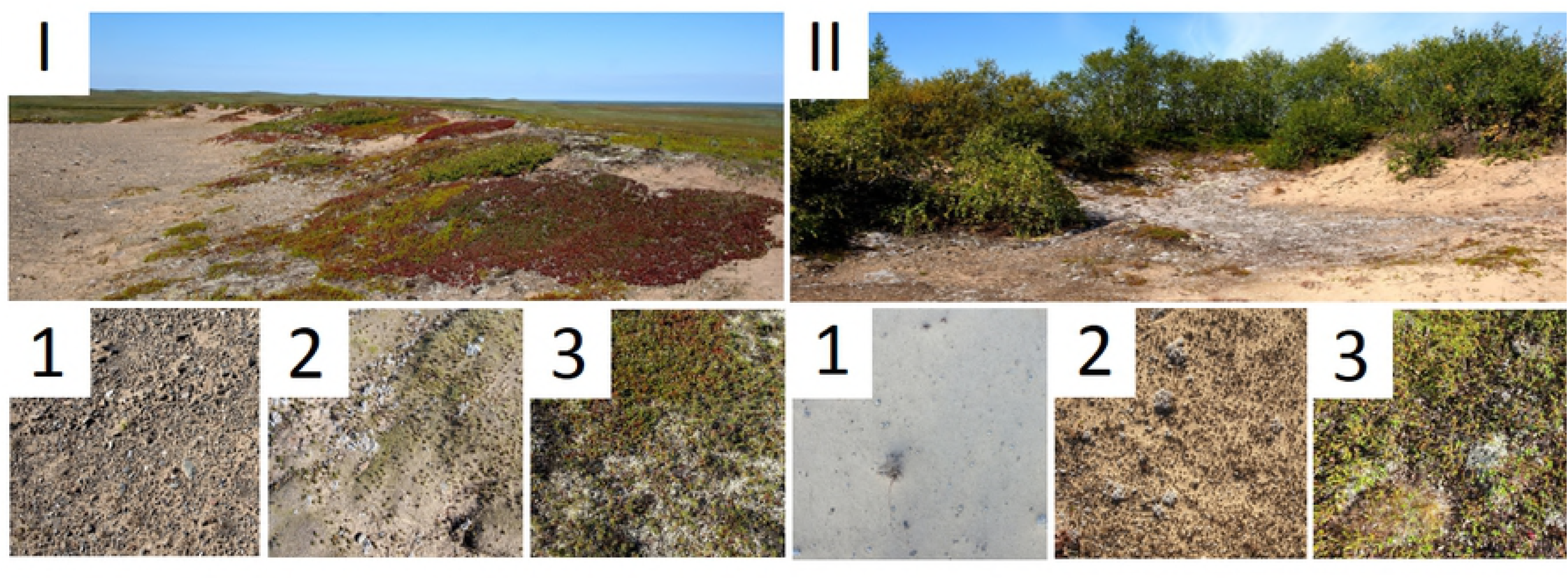
Sampling plots of two chronosequences. Roman numerals indicate sampling site: I – near Nelmin Nos, II – near Naryan-Mar. Indices indicate the type of the plot: US – unfixed sand, SF – semi-fixed surface, MS – mature soil.

The first site (Site I) was located on a flat sand hill, probably a moraine formation, with a height of approximately 30 m (67°58′34.3”N, 52°55′19.9”E, near Nelmin Nos). The areas of unfixed surface (sand with gravel and rare moss and grass shoots) were found on the hilltop. Presumably, the substrate on that surface was unfixed due to wind and snow erosion. The semi-fixed surface was covered by scanty vegetation: the cushion-like moss *Racomitrium canescens*, the lichen *Stereocaulon paschale*, rare subshrubs and other lichens. Vegetation on the mature soil was typical for the tundra zone: lichens (mostly *Cladonia arbuscula* and *Flavocetraria nivalis*), subshrubs (g. *Empetrum*, g. *Arctostaphylos*, g. *Ledum)*, and f *Gramineae.* The soil was classified as Arenosol in the WRB classification [23].

The second site (Site II) was located in a deflation basin with unfixed aeolian sand, which formed small dunes and was gradually covered by vegetation (67°36′23.2”N, 53°08′12.2”E, near Naryan-Mar). The unfixed surface was an aeolian sand without gravel. The semi-fixed surface was partly covered by shoots of moss (g. *Polytrichum*) and lichens (mostly *Stereocaulon paschale*). Vegetation on the mature soil consisted of various lichens, subshrubs (g. *Empetrum*, g. *Arctostaphylos, Vaccinium vitis-idaea*), grasses (*Festuca rubra)* and small trees (g. *Juniperus*, g. *Betula*). The soil was classified as Arenosol in the WRB classification [23] or Psammozem on buried podzol.

For every surface type on each site, five samples of sands and mature soils were taken from depths of 1-5 cm that lacked plants, mosses and lichens. Sampling plots of different types were located on a transect with 3-5 m between each plot. For molecular analyses, samples were stored at −70 °C. The total organic carbon (TOC) and total nitrogen (TN) contents were estimated for the average sample from each plot using a Vario MACRO Cube CN-analyser (Elementar Analysensysteme GmbH, Germany).

### DNA extraction

Total DNA was extracted from 0.5 g of frozen samples using the FastDNA^®^ SPIN kit for Soil (MP Biomedicals, USA) as recommended by the manufacturer. The homogenization step was performed with a Precellys 24 homogenizer (Bertin Technologies, France), program 5 (30 sec, 6500 rev. / min). DNA quality was estimated by electrophoresis in agarose gels (1% w/v in TAE) with further visual DNA detection using the Gel Doc XR+ System (Bio-Rad Laboratories, USA).

### Quantitative PCR analysis

qPCR assays were used for quantitative estimation of ribosomal and N-cycle genes. To estimate the functional potential of microbial communities for N-fixation, ammonia-oxidation and denitrification, the abundance of genes encoding key enzymes of these processes (*nifH, amoA, nirK* and *nirS*, respectively) was measured [18]. 16S ribosomal genes of Bacteria and Archaea, the ITS region of Fungi and functional genes *nifH*, bacterial *amoA, nirK* and *nirS* were quantified using primer sets described in Table 1. All reactions were performed in a C1000 Thermal Cycler with the CFX96 Real-Time System (Bio-Rad Laboratories, USA). The qPCR mix contained 10 μl of 2X concentrated master mix for qPCR (SYBR Green Supermix (Bio-Rad Laboratories, USA) for the ITS region of Fungi, BioMaster HS-qPCR SYBR Blue (Biolabmix, Russia) for the other genes), 0.5-0.8 μM of each primer, and 1 μl of extracted soil DNA template (5-10 ng stock) in a total volume of 20 μl. Quantification of the initial gene copy abundance was performed in CFX Manager. PCR conditions for ribosomal genes were 3 min at 95 °C, followed by 49 cycles of 95 °C for 10 sec, 50 °C for 10 sec, and 72 °C for 20 sec. PCR conditions for N-cycle genes were 3 min at 95 °C, followed by 40 cycles of 95 °C for 20 sec, 54 °C for 20 sec, and 72 °C for 20 sec. To ensure qPCR specificity, melting curve analysis was performed (from 65 °C to 95 °C with an increment of 0.5 °C). Triplicate standard curves ranged from 103 to 108 gene copy number/μl. Standards were made by purifying PCR products and quantifying the concentration by Qubit fluorometer 2 (Thermo Fisher Scientific, USA). Reference organisms for the construction of standard curves for PCR products are described in the Table 1. Efficiencies of qPCR were 82-101% and coefficients of determination were R2 > 0.90 for all standard curves.

**Table 1.**
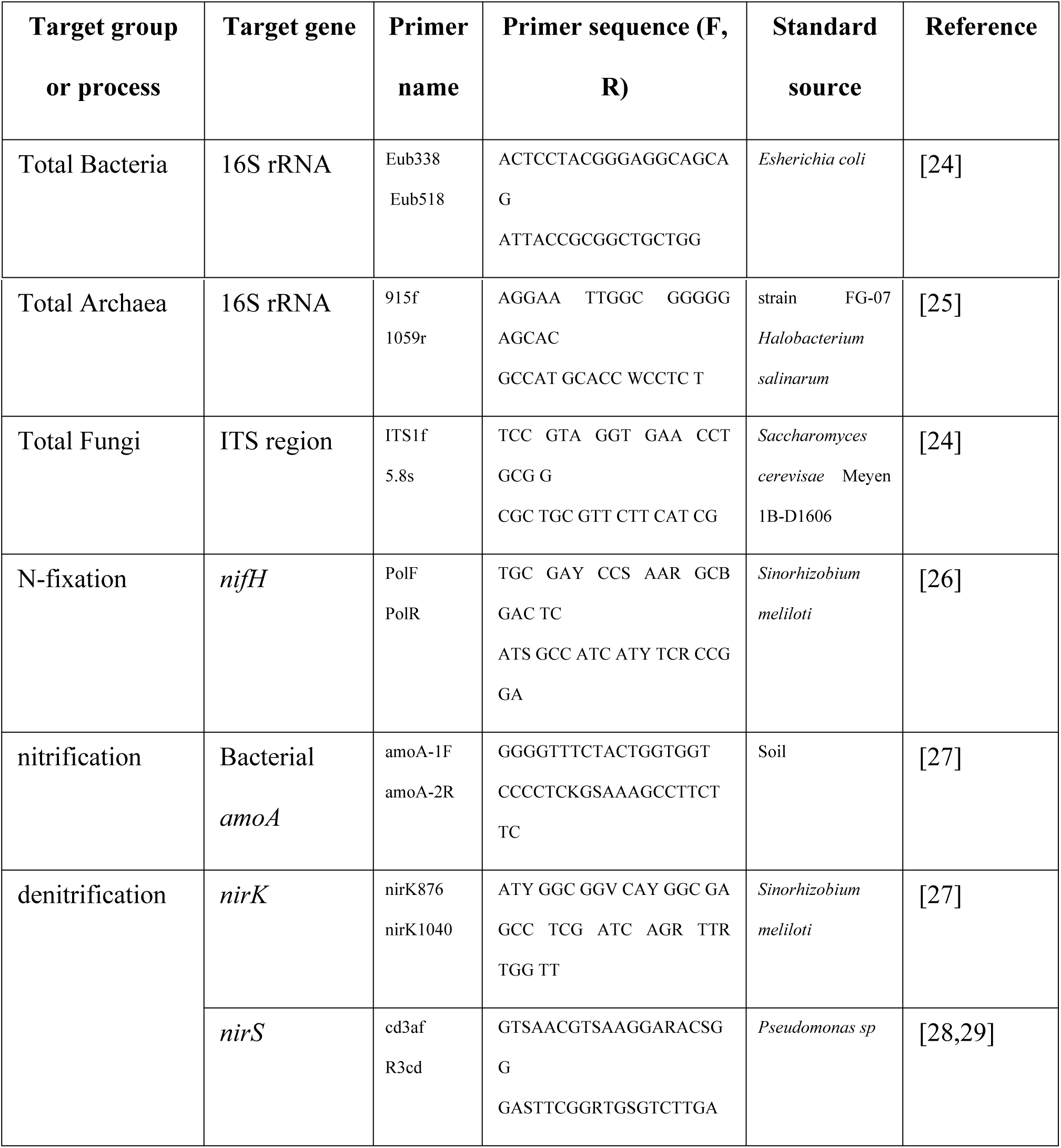
Information about primers and standards for qPCR

### Sequencing of 16S rRNA gene libraries

High-throughput sequencing of the 16S rRNA gene libraries was performed for 5 replicates of each studied sample. The purified DNA isolates were amplified with universal multiplex primers F515 (5′-GTGCCAGCMGCCGCGGTAA-3′) and R806 (5′-GGACTACVSGGGTATCTAAT-3′) [30] targeting variable regions V3-V4 of bacterial and archaeal 16S rRNA genes. PCR was carried out in a 15 μl reaction mixture containing 0.5-1 units of Phusion Hot Start II High-Fidelity polymerase and 1X Phusion buffer (Thermo Fisher Scientific, USA), 5 pM of forward and reverse primers, 10 ng of DNA matrix and 2 nM of each dNTP (Thermo Fisher Scientific, USA). The mixture was denatured at 94 °C for 1 min, followed by 35 cycles of 94 °C for 30 sec, 50 °C for 30 sec, and 72 °C for 30 sec. The final elongation was carried out at 72 °C for 3 min. PCR products were purified according to the recommended Illumina technique using AM Pure XP (Beckman Coulter, USA). Further preparation of the 16S rRNA gene libraries was carried out as described in the MiSeq Reagent Kit Preparation Guide (Illumina, USA). Sequencing of 16S rRNA gene amplicons was carried out on an Illumina MiSeq platform using MiSeq® Reagent Kit v3 (600 cycles) with forward and reverse reading. The raw data is deposited in NCBI database (BioProject ID: PRJNA497067).

### Processing of 16S rRNA gene data

Sequencing data were processed using QIIME [31] and Trimmomatic [32]. Forward and reverse reads that had an overlap of at least 180 nucleotides were merged using the fastq-join algorithm. Sequencing quality tests and operational taxonomic units (OTU) picking based on 97% nucleotide similarity were performed in the QIIME environment. Reference sequences for OTU selection as well as taxonomic affiliation were obtained from SILVA database version 128, 2017 (https://www.arb-silva.de/download/archive/qiime). Singletons (OTUs containing only one sequence) and 16S rRNA sequences of chloroplasts and mitochondria were removed.

### Statistical and sequence analyses

Statistical analysis of gene abundance data was performed in Microsoft Excel and STATISTICA 10.0.

Several indices were used for the estimation of total diversity of the studied prokaryotic communities (α-diversity). The Shannon index was calculated (H *=* Σ *pi lnpi*, where *pi* is the relative abundance of species *i* in the community). The Chao1 index and the phylogenetic diversity whole tree metric were calculated for characterization of the real number of OTUs in the prokaryotic community [33,34]; they were compared with the total number of observed OTUs. Data were normalized to 4090 sequences per sample. A multiple t-test was performed to test for significant (p<0.05) differences of individual microbial taxa and diversity indices.

The analysis of structural differences between prokaryotic communities (β-diversity) was performed using binary metrics of similarity - weighted UniFrac [35]. Based on weighted UniFrac distances, non-metric multidimensional scaling (NDMS) was carried out to construct diagrams of similarity in prokaryotic community structures.

## Results

### Chemical properties of substrates on the studied plots

Both TOC and TN contents increased in correspondence with the stage of soil formation. For both sampling sites, unfixed sands and semi-fixed surfaces had extremely low organic carbon (0-0.18 %) and nitrogen (0.02-0.04 %) contents, while mature soils had much higher amounts of C and N (Table 2). The difference in percentage of TOC between the studied plots was higher than the difference in N content.

**Table 2.**
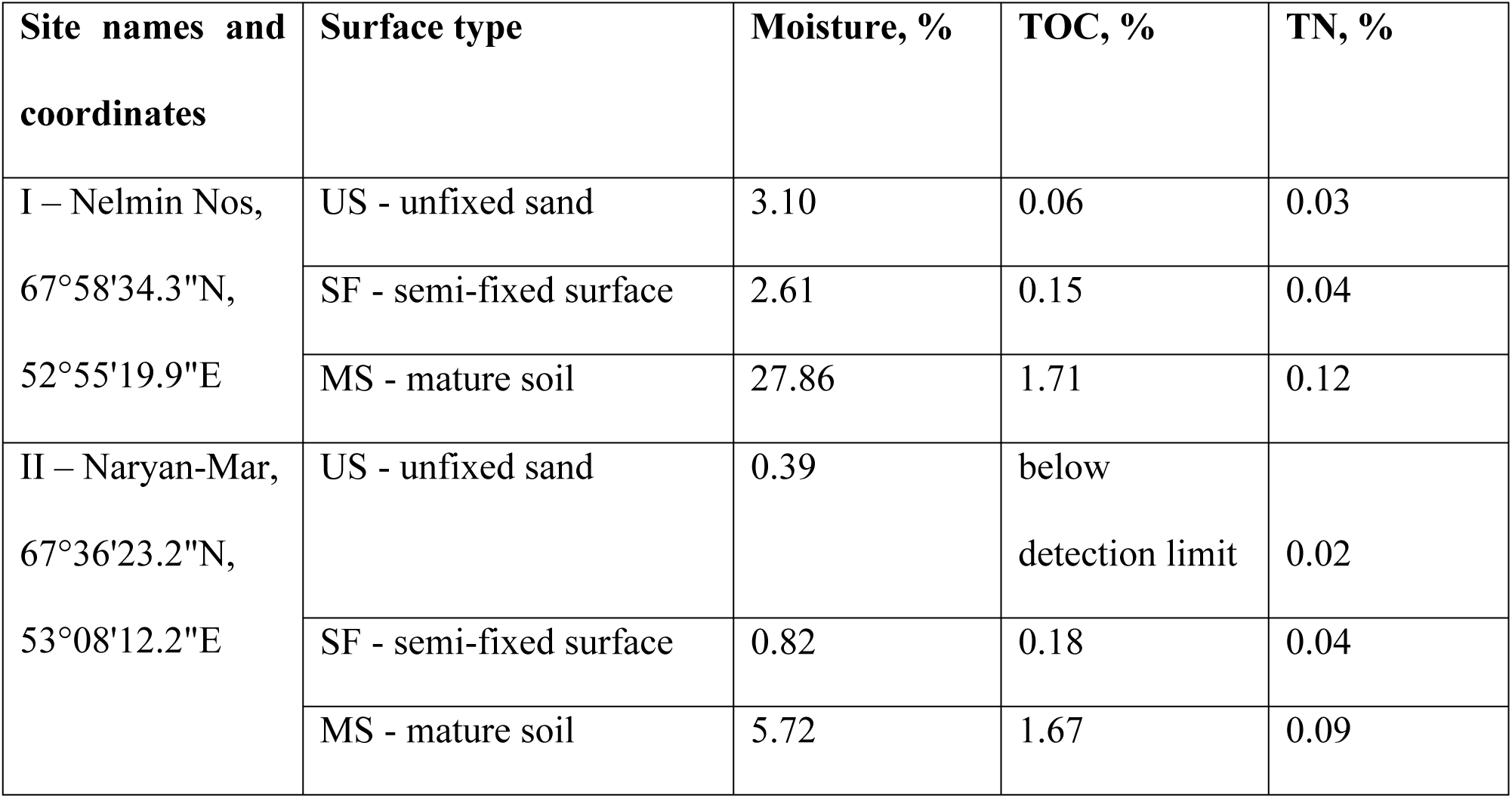
Chemical properties of sampled substrates

### Ribosomal and N-cycle gene abundances in the two chronosequences

The gradual increase in ribosomal gene copy numbers was revealed in both chronosequences from the unfixed sand to the mature soil. At Site I, there was a statistically significant increase (p<0.001) of one order of magnitude in bacterial and fungal gene copy numbers per gram of substrate, while archaeal gene copy number increased by two orders of magnitude (Fig 2). The gene abundances in the samples from the semi-fixed surfaces were intermediate between those of the unfixed sand and the mature soil. An increase of two orders of magnitude in the number of all ribosomal genes was also observed for Site II; however, gene copy numbers in the semi-fixed surface and in the mature soil did not significantly differ from each other (Fig 2).

**Fig. 2.**
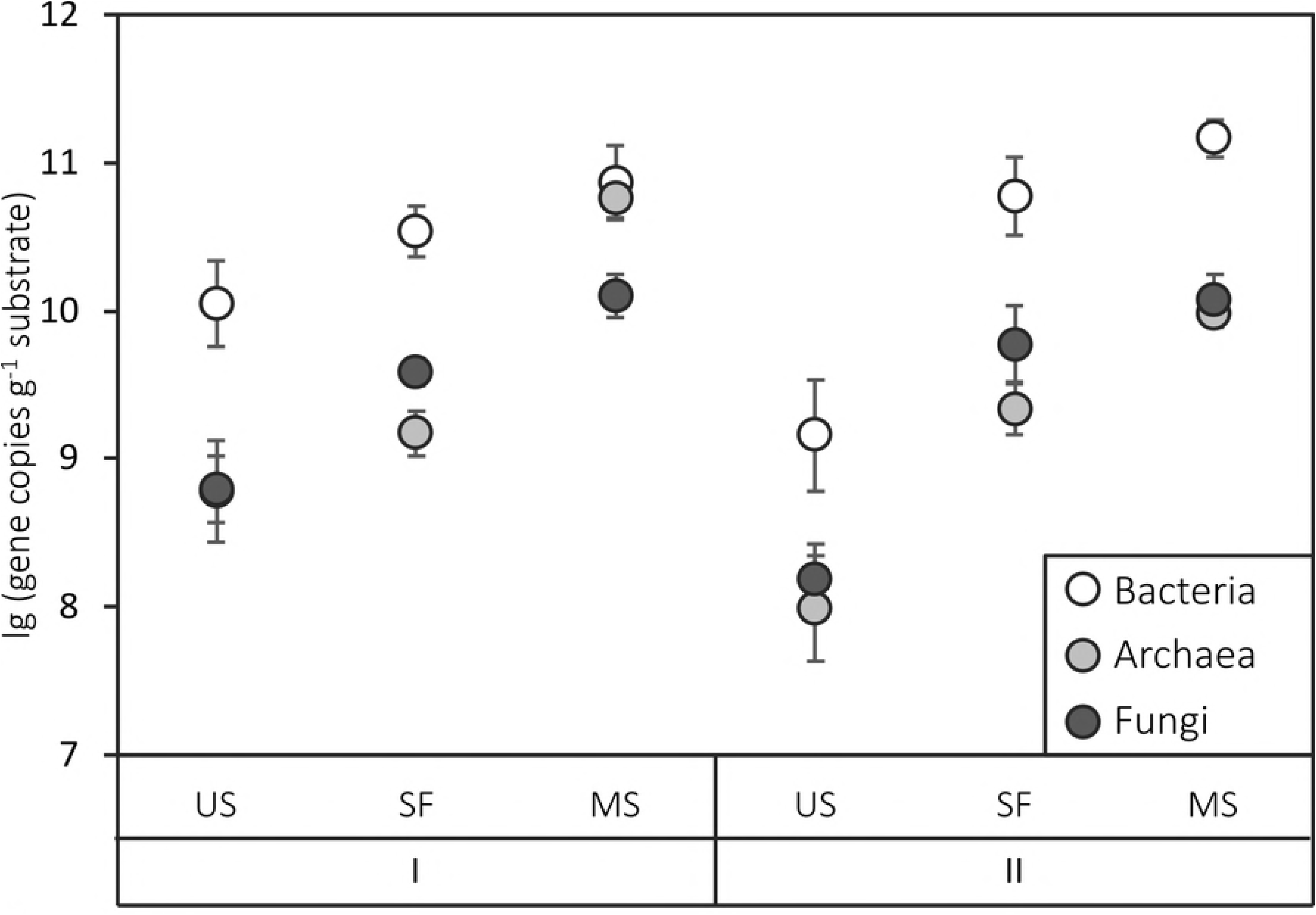
Ribosomal gene copy number in the plots of Sites I and II. US indicates unfixed sand, SF – semi-fixed surface, MS – mature soil. The data are shown as means (n = 5). Error bars represent standard deviations.

Similar trends were observed for the distribution of functional genes along the chronosequences (Fig 3). For both sites, there was a two-order increase in the amount of all N-cycle genes from the unfixed sand to the mature soil. At Site I, the semi-fixed surface was more similar to the unfixed surface; at Site II, conversely, there was stronger similarity between the semi-fixed surface and the mature soil. The *nirK* gene, which is associated with denitrification, was the most abundant among the investigated functional genes in all samples. The number of *nifH* genes (associated with nitrogen fixation) and *nirS* (associated with denitrification) ranged from 4×10^6^ to 7×10^8^ gene copy number g^−1^ substrate. The number of *amoA* genes associated with nitrification was the lowest.

**Fig. 3.**
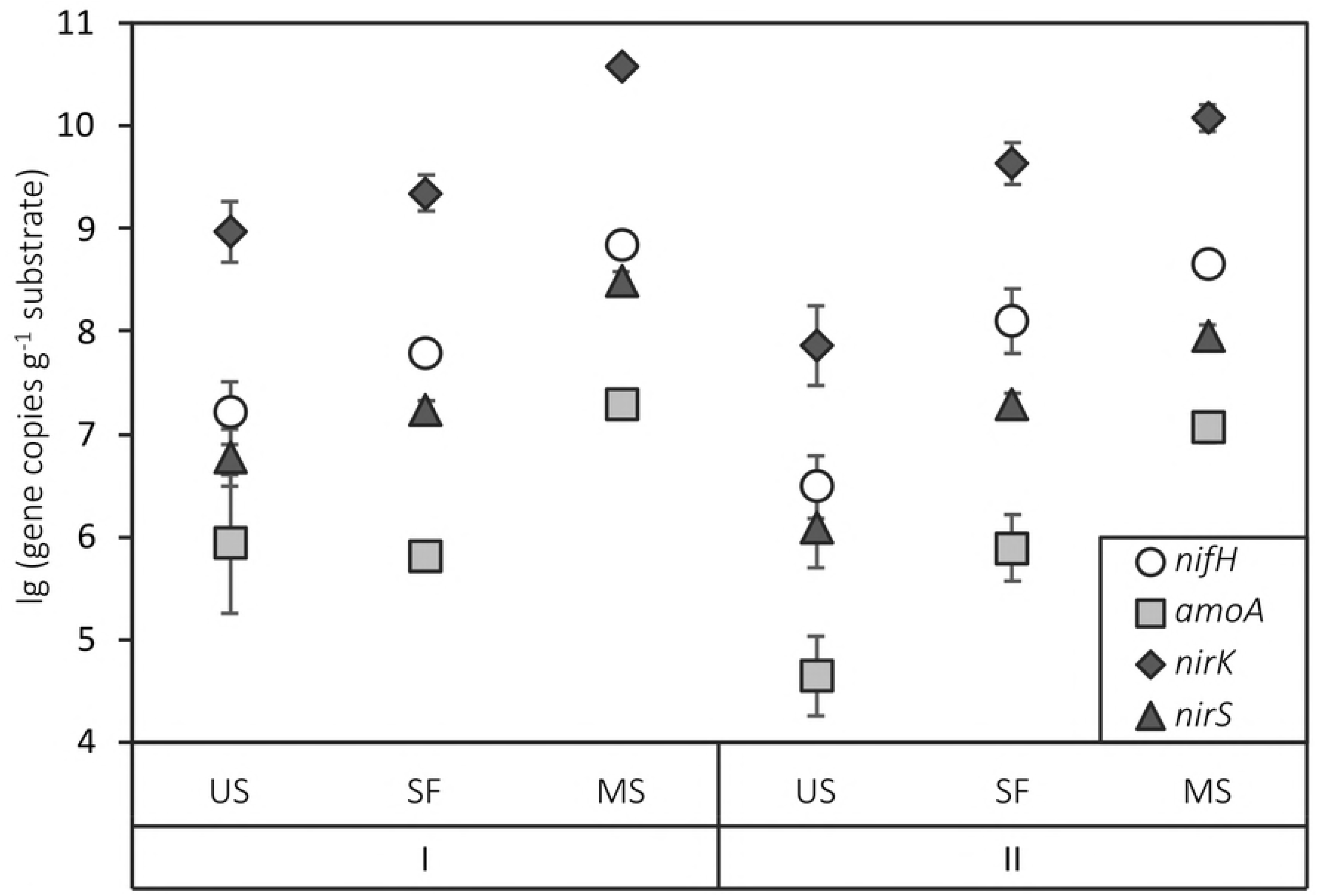
Functional gene copy number in the plots of Sites I and II. US indicates unfixed sand, SF – semi-fixed surface, MS – mature soil. The data are shown as means (n = 5). Error bars represent standard deviations.

Bacterial gene abundance in the substrate correlated with TOC percentage (p<0.05), while archaeal gene abundance correlated with TN (p<0.05), and fungal gene abundance correlated with both TOC and TN (Table 3). All functional genes related to the nitrogen cycle were significantly correlated with TN (p<0.01 for *nirK* and *nirS*, p<0.005 for *nifH* and *amoA*).

**Table 3.**
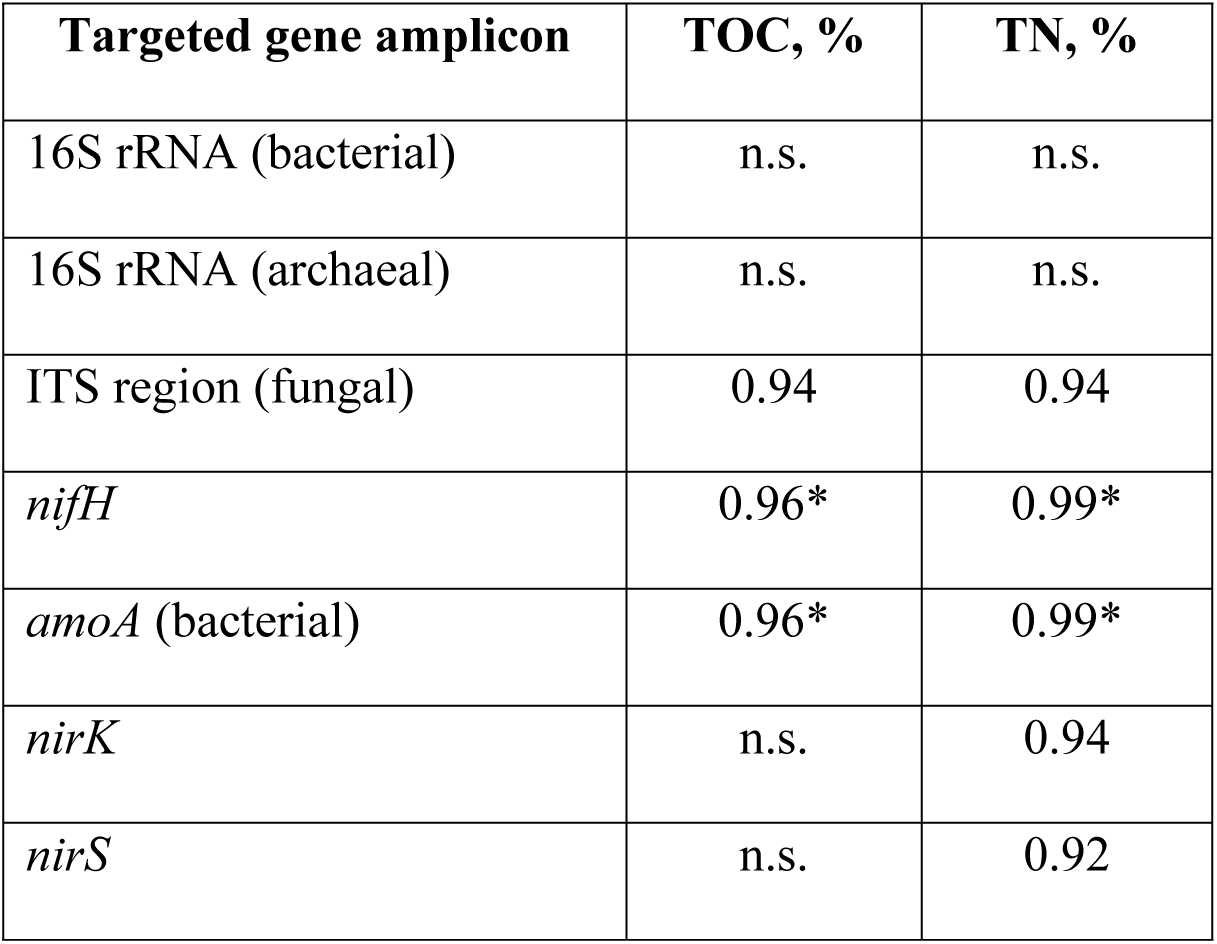
Significant correlations (p<0.01, * - p<0.005) between total organic carbon (TOC), nitrogen (TON) percentage and gene abundances.

### Prokaryotic community structure among the chronosequences

In total, 261 161 sequences of the 16S rRNA gene were obtained (from 2458 to 21 196 sequences per sample) with a mean length of 292 bp.

Phyla *Proteobacteria* and *Acidobacteria* were predominant in all samples (up to 35 % of relative abundance) (Fig 4). The taxonomic structure on the phylum level was similar for the prokaryotic communities in the two mature soils under different vegetation types. The comparison of abundances of different phyla showed that the relatively high abundances of *Thaumarchaeota* (up to 7 % on Site I), *Chloroflexi* (up to 12 %) and *Cyanobacteria* (up to 14 % on Site II) were associated with unfixed sands, while *Planctomycetes* was more abundant in the mature soils and semi-fixed surfaces (Fig 4). The prokaryotic community structure of semi-fixed surfaces was intermediate between those of unfixed sands and mature soils. *Chloroflexi* was relatively more abundant in all samples of unfixed sands and semi-fixed surfaces and contained both cultivable and uncultivable genera, including family *Ktedonobacteraceae.* Genera *Chamaesiphon* (up to 9 % relative abundance), *Crinalium, Leptolyngbya* and *Stigonema* belonging to phylum *Cyanobacteria* were most abundant in the unfixed sand from Site II.

**Fig. 4.**
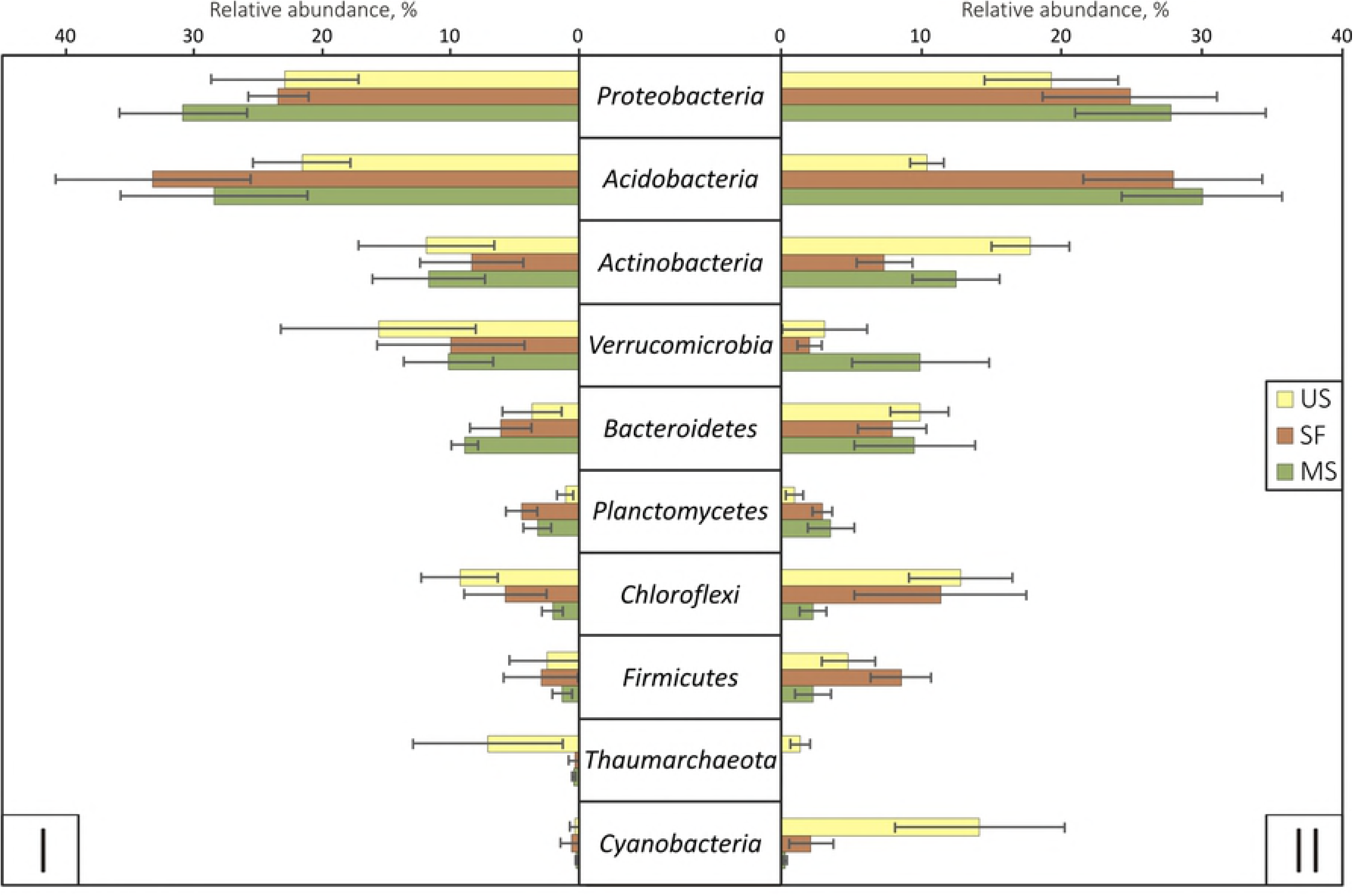
Relative abundances of different phyla in the plots of Sites I and II. US indicates unfixed sand, SF - semi-fixed surface, MS - mature soil. Error bars represent standard deviations.

### The diversity of prokaryotic communities among the chronosequences

The highest prokaryotic α-diversity was found in mature soil from Site I (Table 4). Prokaryotic diversity indices for unfixed sand and the semi-fixed surface at Site I did not differ significantly. Increased α-diversity was observed among the chronosequence at Site I, while no significant difference was revealed for prokaryotic communities at Site II due to high variation between indices for samples from each plot. However, Shannon indices for semi-fixed surfaces of both chronosequences were significantly lower than indices for mature soils and unfixed sands. For all samples, the number of observed OTUs was approximately 65-70 % of the Chao1 index.

**Table 4.**
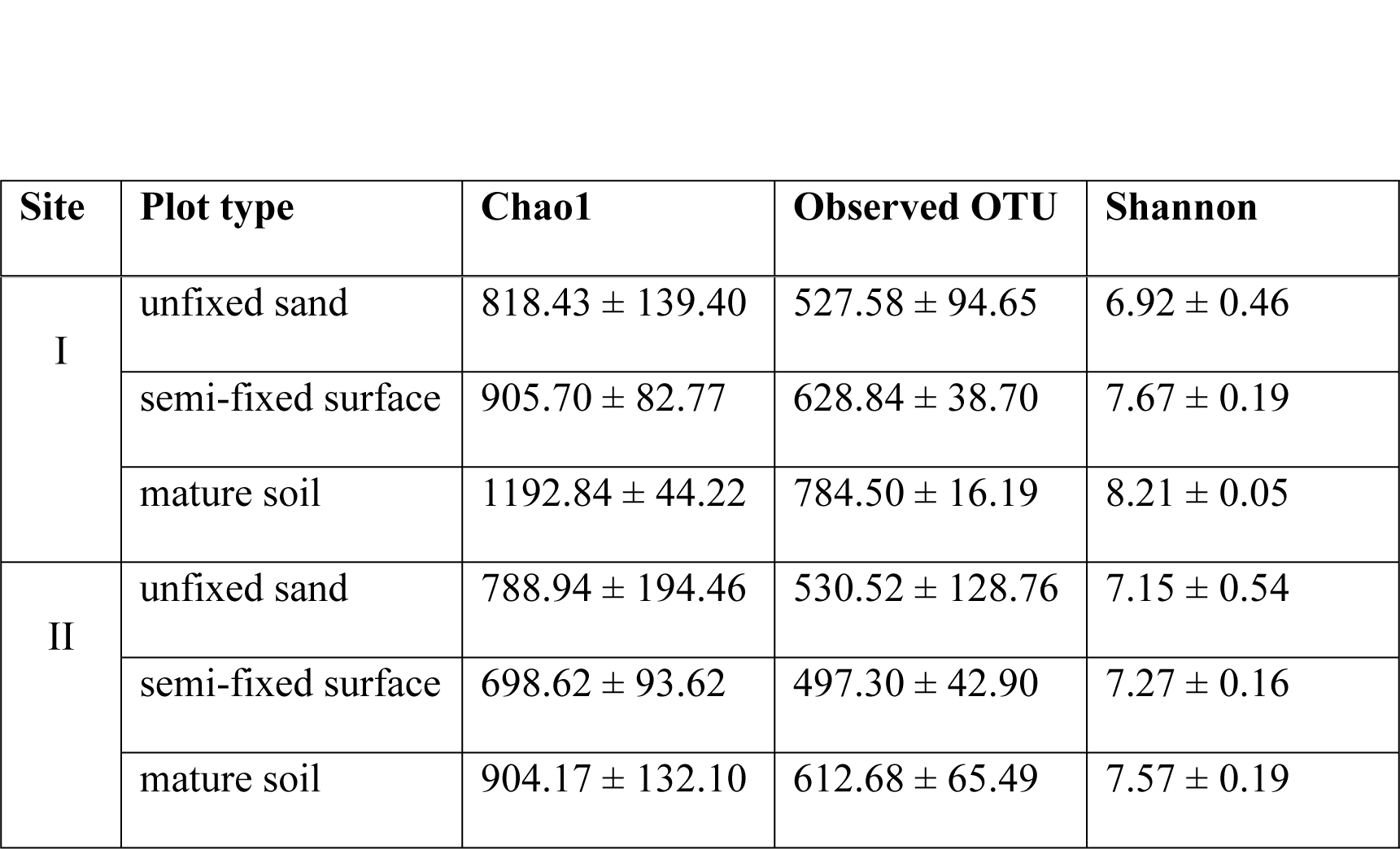
Diversity indices with standard deviation measured for 4090 OTUs for 5 replicates in prokaryotic communities of two chronosequences.

The shift in prokaryotic community composition during the process of sand fixation was observed in both chronosequences (Fig 5). Both unweighted and weighted UniFrac analyses showed similar community compositions of the two mature soils and semi-fixed surfaces, while the sand samples formed separate clusters and differed from each other. In weighted UniFrac metrics, prokaryotic communities of unfixed sands were separated from those of semi-fixed surfaces and mature soils. Bray-Curtis metrics also showed that prokaryotic communities of the two sands were significantly different from the other samples.

**Fig. 5.**
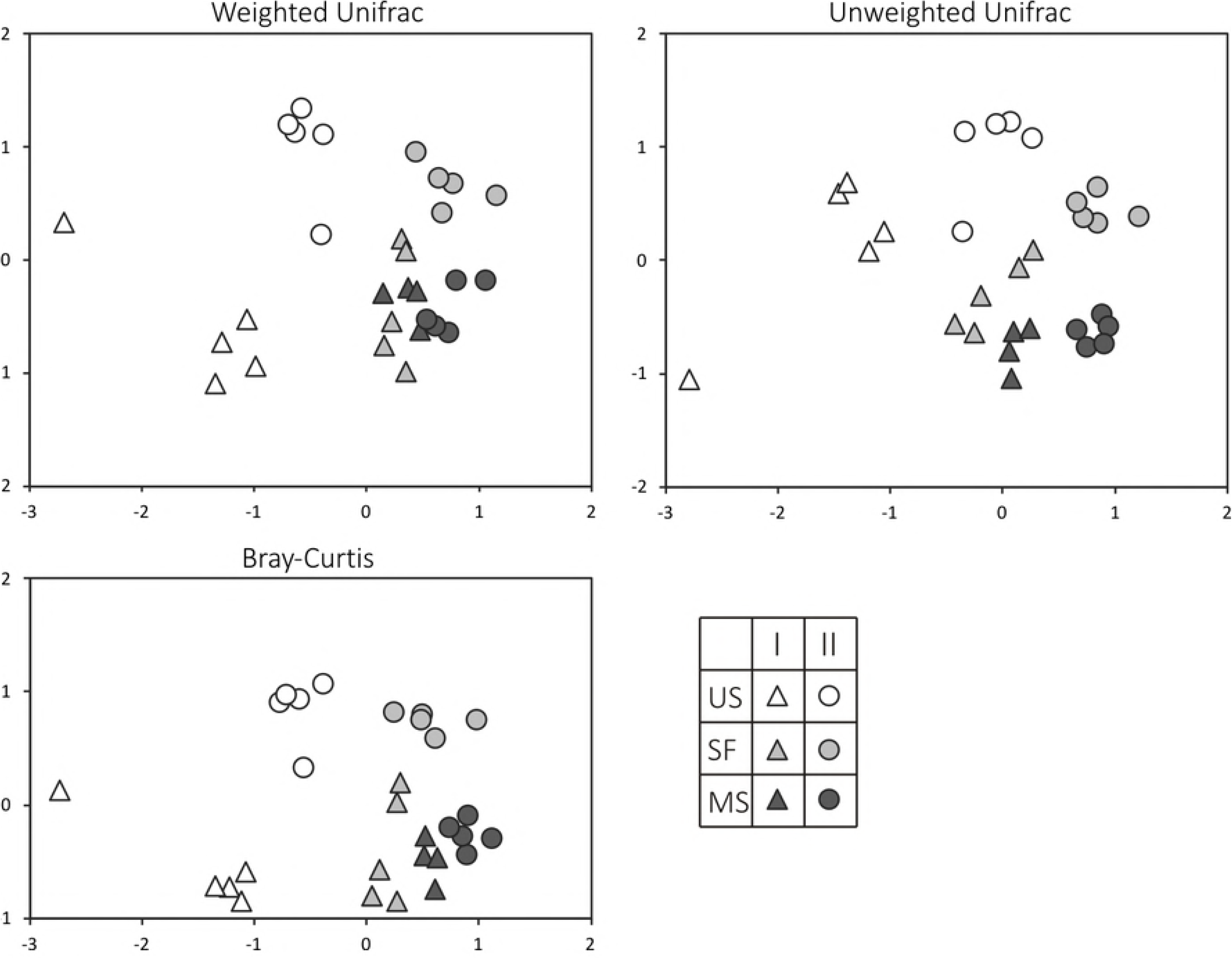
Beta-diversity indices of microbial communities in samples. I, II are the site numbers, US indicates unfixed sand samples, SF – semi-fixed surfaces, MS – mature soil samples.

## Discussion

### Quantitative analysis of ribosomal and N-cycle genes in the two chronosequences

The observed ribosomal gene copy numbers both in semi-fixed surfaces and in mature soils correspond with the previously obtained data on the microbial population abundance for soils and soil-like substrates in northern latitudes [36,37]. The higher abundance of bacterial ribosomal genes in comparison to the abundance of fungal genes in all samples can be explained by low diversity of plant communities at the studied plots; as previously shown in other studies, Fungi are more dependent on the plant communities of barren substrates than Bacteria [21]. However, the abundance of genes increased from unfixed sand to mature soil in both chronosequences, which can be an effect of plant community development and consequently higher available organic carbon and nitrogen contents. Available organic matter in soil is known to be a limiting factor of microbial community development [38], and a correlation between the quantity of ribosomal genes and soil organic carbon content was previously observed for other soils [39].

The abundances of all functional genes associated with transformation of nitrogen also increased in both chronosequences from unfixed sand to mature soil and correlated with total nitrogen content. A relatively high abundance of *nifH* genes related to N_2_ fixation was found in all samples compared to other soils [40,41]. This finding is consistent with the study of metagenomes of Arctic tundra soil, where N-assimilation genes were present in all bacterial genomes in microbiomes of different types of polygonal landscapes [42]. Another study showed that in soils formed on glacial retreats, the nitrogen fixation rates significantly increased during the first 4-5 years of succession [12]. Thus, nitrogen-fixing bacteria are important for soil microbial community assemblage and functioning, especially in the tundra zone. The lowest abundance of the bacterial *amoA* gene among all functional genes studied can be explained by the low amount of organic nitrogen in all samples. Bacterial *amoA* gene abundance in soil is known to be related to the available ammonia concentration [43]. In all samples, *nirK* gene abundance was the highest in comparison to other N-cycle genes. Genes associated with denitrification (*nirK* and *nirS*) were previously found to be more abundant than *nifH* and *amoA* in other soils [40]. The abundances of the two nitrite reductase-encoding genes (*nirK* and *nirS*) observed in this study were disproportionate. According to previous studies, *nirK* (copper nitrite reductase) is more widespread in terrestrial ecosystems, while *nirS* (cytochrome *cd1* nitrite reductase) is more abundant in marine environments [44]. Moreover, *nirK* genes were more abundant than *nirS* genes in Arctic soils [45]. *nirS* is more widely distributed among denitrifiers, but *nirK*-containing denitrifiers are more physiologically variable [46]. However, in some soils *nirS* genes were more abundant than *nirK* genes [47,48].

Thus, the obtained data for both Sites supported the hypothesis of increased gene abundance among the chronosequences. The similar dynamics of ribosomal and functional gene abundance in the two chronosequences can be explained by the congruent patterns of increased total organic carbon and total nitrogen from unfixed sands to mature soils on the two sites. Gene copy number indirectly indicates the biomass of different functional and taxonomic groups in soil microbial communities, which increases during primary succession on different types of barren substrates [7].

### Changes of prokaryotic community structure during soil formation

The prokaryotic communities of all samples were dominated by phyla *Acidobacteria* and *Proteobacteria*, which was previously observed for other soils of the tundra zone [42,49].

Phylum *Thaumarchaeota* was found to be relatively more abundant in unfixed sands of both sites. All OTUs belonging to *Thaumarchaeota* were uncultivable Archaea and were previously observed in other terrestrial environments. Among all Archaeal phyla, *Thaumarchaeota* are known to be predominate in Arctic and Antarctic soils [49]. We suggest that the relative abundance of *Thaumarchaeota*, but not their absolute number, decreased from unfixed sands to mature soils because archaeal gene abundances in the unfixed sands were a hundred-fold lower than in the mature soils.

Family *Ktedonobacteraceae* belonging to phylum *Chloroflexi* that were predominate in unfixed sands are known to be negatively correlated with organic matter content in deforested soil [50]. This family is mostly represented by uncultivable genera; its cultivable representatives are filamentous, aerobic and mesophilic [51]. *Ktedonobacteraceae* was previously found in young soils [20,52,53]; thus, we suggest that representatives of this family are important at initial successional stages.

Representatives of phylum *Cyanobacteria* were more abundant in the samples of unfixed sand from Site II, which can be explained by low levels of insolation of sand on Site I due to gravel cover. *Cyanobacteria* are known as free-living phototrophs capable of nitrogen fixation, especially in extreme environments [13,49,54]. Representatives of this phylum could be the primary producers of organic matter in unfixed sands due to the lack of organic carbon and nitrogen. A decrease in *Cyanobacteria* abundance and number of observed OTUs belonging to this phylum with soil age was previously observed for soils formed by glacial isostatic adjustment in Fennoscandia [55]. Genus *Leptolyngbya* was found on plots of unfixed sand on Site II. It is known as a producer of adhesive extra-cellular polysaccharides and organic acids that can degrade rock [56]. Thus, all these taxa inhabit barren substrates, and their active presence in the community can be considered an indicator of the primary stage of development of microbial succession.

### Diversity changes among the chronosequences

The gradual increase of prokaryotic α-diversity from initial stages of sand fixation to mature soils was expected for both chronosequences. However, prokaryotic α-diversity increased from the unfixed sands to the mature soil on Site I, while prokaryotic communities of all samples on Site II did not follow the same pattern, and their α-diversities did not change with the successional stage. The obtained results somewhat contradict previously discovered trends of incremental growth of prokaryotic α-diversity during revegetation on moving dunes [10]. However, some studies reported the highest prokaryotic diversity at the early stages of soil formation [57,58]. Although the microbial biomass on barren substrates was relatively low, the high diversity of the unfixed sand prokaryotic community can be explained by the variety of necessary adaptations to harsh environmental conditions.

The prokaryotic community structures in samples of the two mature soils appeared to be very similar to those in the unfixed sand samples. The estimation of β-diversity showed the same pattern of unevenness of prokaryotic communities in the unfixed sand samples. This observed dissimilarity could be a consequence of random propagule input in unfixed sands, while the developed vegetation on both mature soils allowed the formation of more stable prokaryotic communities.

## Conclusions

Using a chronosequence approach, we found the expected trends in microbial populations: increased microbial community abundance and change of prokaryotic community structure from unfixed sands to mature soils. The highest prokaryotic diversity and abundance, as well as the amount of microorganisms involved in the nitrogen cycle, were revealed in mature soil under developed plant cover. However, the prokaryotic diversity during soil formation increased slightly, with the minimum values found in sand under pioneer vegetation (intermediate stages of succession). In contrast with our predictions, prokaryotic communities under the two unfixed sands were highly dissimilar, while different plant cover on the two mature soils shifted microbial community structure in the same way and reduced spatial heterogeneity.

## Acknowledgments

We would like to thank Sergei Khokhlov and Dmitry Konuyshkov for their support. Experiments were performed using the equipment of the Collective Use Center “Functions and Properties of Soils and Soil Cover” of the Dokuchaev Institute. High-throughput sequencing was performed using the Illumina MiSeq platform of the Common Use Center of Saint Petersburg State University.

